# Optimal pathways control fixation of multiple mutations during cancer initiation

**DOI:** 10.1101/2022.02.07.479413

**Authors:** Hamid Teimouri, Cade Spaulding, Anatoly B. Kolomeisky

## Abstract

Cancer starts after initially healthy tissue cells accumulate several specific mutations or other genetic alterations. The dynamics of tumor formation is a very complex phenomenon due to multiple involved biochemical and biophysical processes. It leads to a very large number of possible pathways on the road to final fixation of all mutations that marks the beginning of the cancer, complicating the understanding of microscopic mechanisms of tumor formation. We present a new theoretical framework of analyzing the cancer initiation dynamics by exploring the properties of effective free-energy landscape of the process. It is argued that although there are many possible pathways for the fixation of all mutations in the system, there are only few dominating pathways on the road to tumor formation. The theoretical approach is explicitly tested in the system with only two mutations using analytical calculations and Monte Carlo computer simulations. Excellent agreement with theoretical predictions is found for a large range of parameters, supporting our hypothesis and allowing us to better understand the mechanisms of cancer initiation. Our theoretical approach clarifies some important aspects of microscopic processes that lead to tumor formation.

## 1 Introduction

Cancer is a collection of genetically related diseases that exhibit uncontrolled cellular growth in specific tissues that strongly affects the normal functioning of living organisms and might even lead to a death [19, 35]. It is now well established that the cause of cancer is the accumulation of several specific genetic or epigenetic alterations in genome [9, 19, 22, 35]. The genetic errors frequently appear during cell replications, but a huge majority of them are quickly repaired by specific biochemical DNA repair pathways [19, 33, 34]. Due to the stochastic nature of these chemical processes, a very small fraction of these mutations might escape the repair mechanisms, and the accumulation of several such mutations eventually can lead to a cancer. However, the microscopic details of how the tumors form still remain not well understood despite significant efforts in multiple research areas [4, 10].

It is clear that tumor formation is the result of complex network of chemical, biological and physical processes [17, 35]. Each type of the cancer is initiated by acquisition of relatively small number of specific mutations (typically between two and eight) [1, 13, 14]. However, the specific sequence of events that lead to cancer is still not well understood [5, 21, 22]. In healthy tissue cells, there are multiple stochastic events of cell replications, cell deaths, and cell mutations that eventually cause cancer. Most probably, the appearance of some mutations are taking place independently from each other. This means that the overall fixation of mutations, i.e., filling the tissue with all mutated cells, which is frequently assumed as the beginning of the cancer, is not a linear process of sequential fixations of each mutation one by one. Indeed, there are many other pathways beyond the sequential one, and the overall phenomenon is known as a stochastic tunneling in the fixation dynamics of several mutations [2, 5, 8, 11]. This complex picture of pre-cancer processes has been supported by some recent experimental observations, including single cell DNA sequencing, multi-region sequencing and deep sequencing [3].

In recent years, several theoretical approaches to analyze the dynamics of cancer initiation have been proposed [5, 12, 16, 25, 31, 32]. Many of them specifically addressed the effect of stochastic tunneling [2, 5, 8, 27], showing the importance of this phenomenon. For example, it was found that stochastic tunneling is a leading process in adaptation dynamics of asexual populations across fitness valleys [7, 36, 37]. However, despite significant advances in our understanding of the cancer initiation dynamics, the main difficulty is that the overall process of multiple mutations fixation has a very large number of possible pathways. This leads to the use of several approximations that are not always able to realistically describe the processes that lead to tumor formation.

In this paper, we present a new theoretical approach of analyzing the cancer initiation dynamics. It is based on analogy with the analysis of complex processes in physics and chemistry, suggesting to explore the effective free-energy description of the fixation dynamics for several mutations. The method is explicitly applied for the system with two mutations where it is tested with analytical calculations and Monte Carlo computer simulations. It is found that there are only few pathways that dominate the fixation process. This significantly simplifies the overall analysis and it allows to understand better the underlying microscopic mechanisms of tumor formation. The extension of these theoretical method to systems with multiple mutations is also discussed. It is interesting to note that our idea is related to recently proposed method of analyzing stochastic systems that takes into account their topological properties [30].

## 2 Methods

The cancer initiation involves various processes such as the appearance of mutations, multiple cell divisions and cell deaths [28]. While the system starts with one of the normal cells becoming mutated, the tumor formation is associated with the situation when all cells in the tissue acquire all mutations, which is known as a mutation fixation. The main idea of our method is to look at the cancer initiation process as a motion of a “particle” in an effective free-energy landscape created by transitions between different stochastic states. The “particle” corresponds to a current state of the system, which is determined by the number of cells with different degrees of mutations. The effective free-energy landscape is built in the following way. Since all transition rates out of the given state can be specified, one can evaluate the residence times in each state. Then the inverse of the residence time correlates with effective free-energy for this state. This is because the longer the system is occupying the given state the lower is the corresponding “free energy”. Finally, the cancer initiation process is viewed as a motion in this effective potential between initial and end states. The initial state has one mutated cell with only one mutation, and the final state has all cells with all possible mutations. The analysis of stochastic dynamics in free-energy landscapes has been widely explored for understanding the mechanisms of various complex processes in chemistry, physics and biology [15, 26].

Let us explain in more detail the hypothesis by concentrating on a simplest non-trivial “two-hit” model of cancer initiation dynamics [13, 14]. In this model, which is schematically shown in Fig. 1a, the system might be without any mutations in the tissue cells (indicated by the green circle), or it might sequentially acquire one mutation in some of the cells (indicated by the yellow circle) and it might finally get two mutations in all tissue cells (indicated by red circle). Two specific examples of the processes that might follow the “two-hit” model, inactivation of tumor suppressor genes and chromosomal instability, are presented in Figs. 1b and 1c [22].

**Figure 1.**
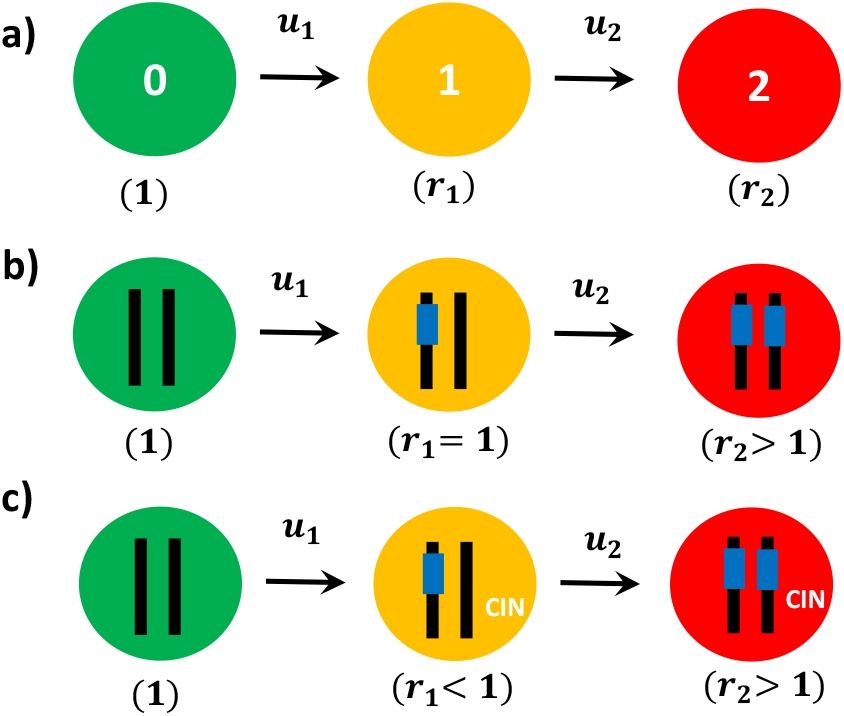
a) A schematic description for cancer initiation dynamics in the ”two-hit” model. First and second mutations can appear with rates *u*_1_ and *u*_2_, respectively. The fitness parameters for normal, one-mutation and two-mutation cells are 1, *r*_1_ and *r*_2_, respectively. b) A schematic view for the process of inactivation of tumor suppressor genes that might follow the ”two-hit” model. In this case, introducing first mutation does not affect the fitness parameter (*r*_1_ = 1), while acquiring the second mutation would increase the cell proliferation rate (*r*_2_ *>* 1). c) A schematic view of the chromosomal instability (CIN) process that also might follow the ”two-hit” model. [23]. Here, the first mutation by itself is known to be disadvantageous (*r*_1_ *<* 1) while introducing the second mutation would make their combination advantageous (*r*_2_ *>* 1).

The total number of cells in the tissue is assumed to be equal to *N* and it is constant during the cancer initiation process [22]. This is one of the features of homeostasis that keeps the number of cells in healthy adult tissues to be constant [22]. As illustrated in Fig. 1a, there are three types of tissue cells that can be found in the system: normal cells without mutations (type 0), cells with one mutations (type 1) and cells with two mutations (type 2). We define *n*_0_, *n*_1_ and *n*_2_ as the number of type 0, type 1 and type 2 cells, respectively, which means that

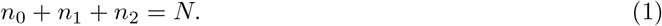

At some time (assumed to be *t* = 0), one of the tissue cells gets a first mutation with a rate *u*_1_. Some time later, a second mutation appears in one of the single-mutated cells with a rate *u*_2_. Note that this might happen before all the tissue cells will have the single mutation, i.e., before the fixation of the first mutation. It is realistic also to assume that the system is in a so-called strong-selection and weak-mutation regime with *u*_1_, *u*_2_ ≪ 1/*N* [6, 20, 22, 24]. This is because the mutations are rare events due to a very low probability of escaping the DNA repair mechanisms. This also means that the dominating processes that evolve the system are cell divisions and removals that at the same time must also keep the total number of cells in the tissue fixed. We define cell division rates for the normal, single-mutated and double-mutated cells as *b, br*_1_ and *br*_2_, respectively. The parameter *r*_1_ is known as a fitness parameter, and it describes how faster the single-mutated cells replicate in comparison with the normal cells. It reflects the overall physiological impact of this mutation on cellular metabolism: if *r*_1_ < 1, the mutation is disadvantageous, *r*_1_ = 1 corresponds to a neutral effect, while for *r*_1_ > 1 the mutation is advantageous. Similarly, the parameter *r*_2_ is a fitness parameter for double-mutated cells that reflects the cumulative effect of both mutations influencing the cellar metabolic processes [32].

Because the total number of cells in the tissue is constant and equal to *N*, there are (*N* + 1)(*N* + 2)*/*2 possible discrete states in the system, as shown in Fig. 2 (left). Each state can be characterized by just two integer numbers (*n*_1_, *n*_2_) with 1 ≤ *n*_1_, *n*_2_ ≤ *N*, which corresponds to the system having *n*_1_ cells with one mutation, *n*_2_ cells with two mutations and *N* − *n*_1_ − *n*_2_ cells without mutations. For example, the state (1, 0) describes the state with one single-mutated cell, zero cells with two mutations and *N* − 1 normal cells without mutations. The state (*N*, 0) describes the situation when all cells have the single mutation, i.e., this is the fixation of the first mutation. In the state (0, *N*) all cells are double-mutated, and this is the fixation of both mutations, which is also assumed to be the beginning of cancer. In our model, the cancer initiation process starts in the state (1, 0) and ends in the state (0, *N*): see Fig. 2.

**Figure 2.**
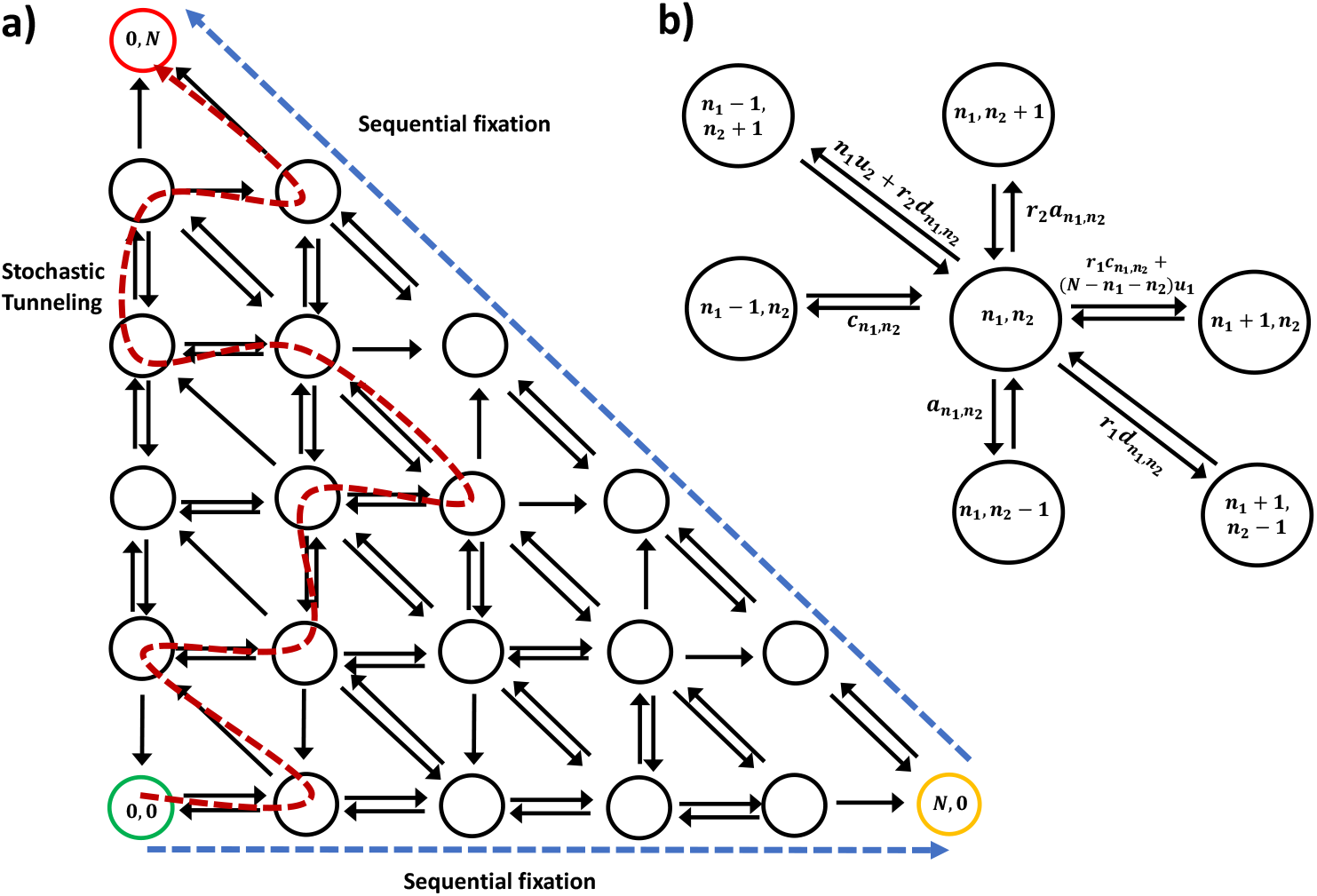
a) A schematic view of the two-dimensional state space associated to fixation of two mutations. Sequential fixation pathway is shown with blue dashed lines, while a possible stochastic tunneling is shown with red dashed line. b) The corresponding stochastic transition in the system. Each state *n*_1_, *n*_2_ corresponds to a population with *n*_1_ cells of type 1 and *n*_2_ cells of type 2.

To construct the effective free-energy landscape for the cancer initiation process in the two-hit model, we need to specify the rates of all possible transitions between different states. As shown in Fig. 2a, there are two types of discrete states in the system depending on the location in the discrete-state stochastic scheme that we might label as bulk and boundary states. It can be shown that the number of bulk states is equal to (*N* − 1)(*N* − 2)*/*2 and there are six possible stochastic transitions out of each of them (see Fig. 2b). The horizontal forward transition rate from the state (*n*_1_, *n*_2_) to the state (*n*_1_ + 1, *n*_2_) is given by the rate 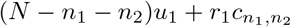 (Fig 2b). The first term in this rate reflects the possibility of acquiring the first mutation by any of the normal cells (*N* − *n*_1_− *n*_2_ of those), while the second term reflects the cell replication by the mutated cell and the removal of the normal cell. The horizontal backward transition rate from the state (*n*_1_, *n*_2_) to the state (*n*_1_ − 1, *n*_2_) is given by 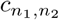, reflecting the cell replication of the normal cell and the removal of the single-mutated cell. Similarly, as shown in Fig. 2b, the forward vertical transition rate from the state (*n*_1_, *n*_2_) to the state (*n*_1_, *n*_2_ + 1) is given by 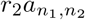 and the vertical backward transition rate is 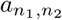. In these cases, the changes in the number of double-mutated cells is taking place only via the cell divisions of the double-mutated cells and removal of the normal cells. More complex processes are taking place in diagonal transitions (see Fig. 2b). The forward diagonal transition rate to go from the state (*n*_1_, *n*_2_) to the state (*n*_1_ − 1, *n*_2_ + 1) is equal to 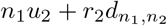. Again, the first term reflects the possibility of the single-mutated cells to acquire the second mutation (*n*_1_ single-mutated cells in the system in the starting state), and the second term is due to the cell replication of the double-mutated cell and the removal of the single-mutated cell. Then the backward diagonal transition from the state (*n*_1_, *n*_2_) to the state (*n*_1_ + 1, *n*_2_ − 1) is given by 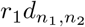, which corresponds to the cell replication of the single-mutated cell and the removal of the double-mutated cell. In the SI, we explicitly show how to evaluate the transition rates 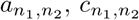 and 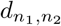, yielding the following expressions

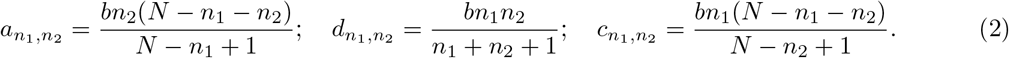

As one can see from Fig. 2a, there are also 3*N* boundary states that lie on the horizontal, vertical and diagonal axis. They correspond to the situation with *n*_0_ = 0 (only single or double-mutated cells, diagonal line) or *n*_1_ = 0 (only normal or double-mutated cells, vertical line) or *n*_2_ = 0 (only normal or single-mutated cells, horizontal line). The number of stochastic transitions out of these states varies between 1 and 3, which is less than 6 due to the boundary location, but the specific rates follow the same rules as for the bulk states.

Knowing all the transition rates between different states allows us to evaluate the average residence times in each state. For any bulk state (*n*_1_, *n*_2_) we have

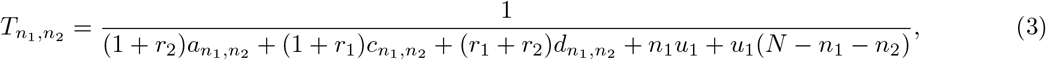

while similar expressions can be also obtained for the boundary states. Then the effective free-energy potential for the system can be estimated via the inverse residence times as

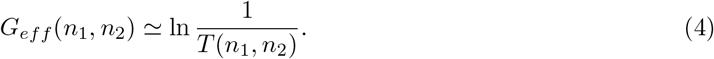

This can be justified by noting that the residence times in each state might be viewed as proportional to the probability for the system to be found in the given state, exp[− *G*_*eff*_ */k*_*B*_*T*]. The results of our estimates for the effective free-energy landscape are presented in Fig. 3a. One can see that there is a broad effective free-energy mountain that correspond to the most bulk states surrounded by free-energy valleys that correspond to the boundary states.

**Figure 3.**
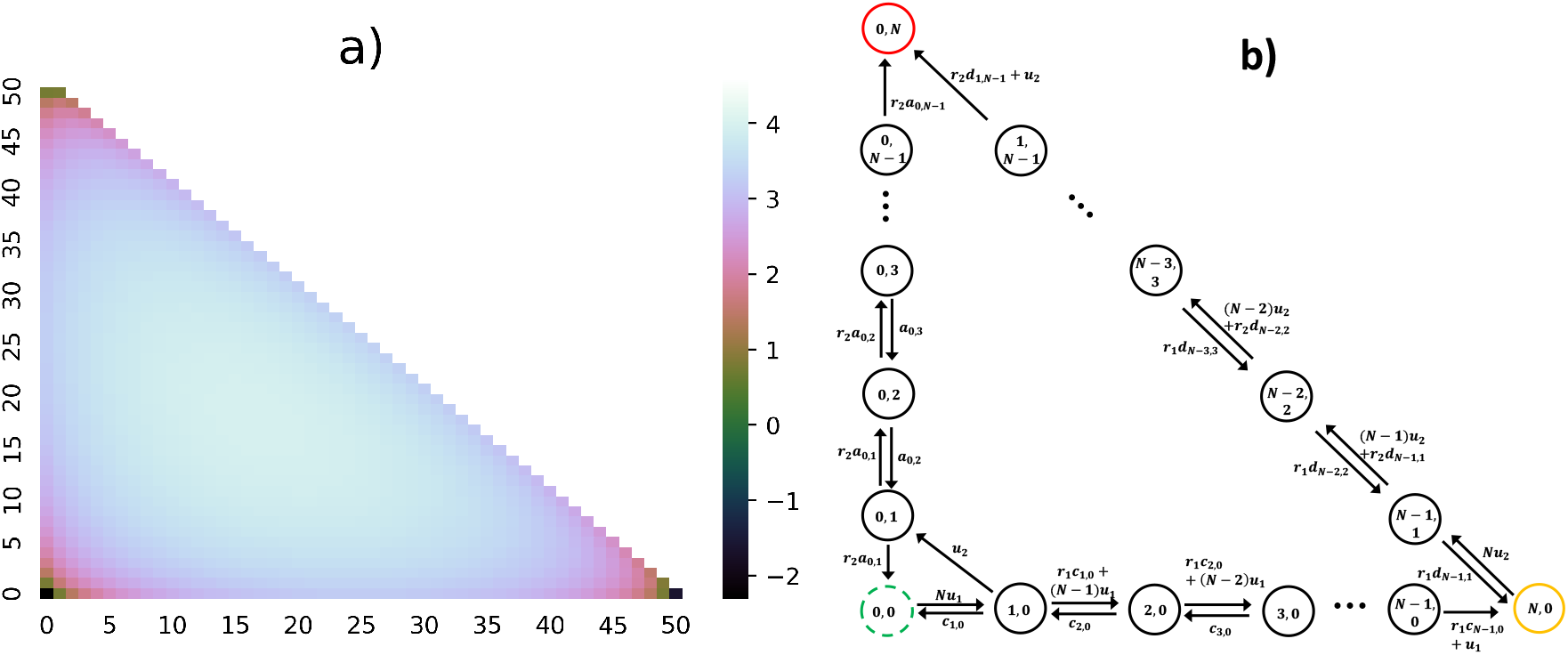
a) The effective free-energy potential that is proportional to inverse residence times in each state, *G*_*eff*_ (*n*_1_, *n*_2_) ≃ 1*/T* (*n*_1_, *n*_2_. In calculations 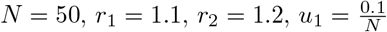 and 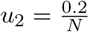 were utilized. b) Schematic view of the optimal pathways that include only the boundary states.

The cancer initiation dynamics in the two-hit model can be investigated by analyzing stochastic transitions in the effective discrete-state scheme presented in Fig. 2a. There are several ways it can be done. In this work, we employ the method of first-passage probabilities that was already successfully applied in earlier studies of cancer initiation dynamics [31, 32]. In this approach, one could define functions *F* (*n*_1_, *n*_2_; *t*) as the probability to reach the final fixation state (0, *N*) at time *t* given that system starts at state (*n*_1_, *n*_2_) at time *t* = 0. The temporal evolution of these functions is described by a set of following backward master equations,

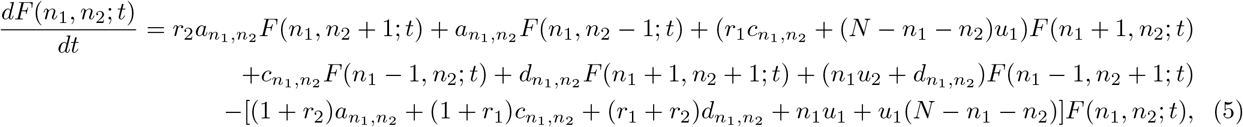

for the bulk states. Similar, but simpler expressions can be obtained for the boundary states. In addition, we have an initial condition *F* (0, *N* ; *t*) = *δ*(*t*), which means that the fixation is instantaneous if the system already starts in the final state. The first-passage probability functions should provide a full description of cancer initiation process. We are particularly interested in the mean fixation probabilities from the state 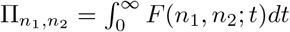 and the mean fixation times 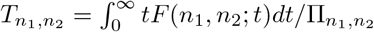, starting from the state (1, 0).

This first-passage approach allows us to obtain exact results but only small values of *N*. In the SI, we provide detailed derivations (supported by Monte Carlo computer simulations) for *N* = 2 and *N* = 3 systems. However, the typical number of tissues cells is much larger, and it ranges from *N* ≃10^5^ to *N* ≃10^9^, but at these conditions it is not feasible to get explicit solutions. It is possible, however, to investigate the cancer initiation dynamics in the system using Monte Carlo computer simulations, but even this approach is challenging because the total number of states is of the order of ≃*N* ^2^.

The analysis of the free-energy landscape presented in Fig. 3a suggests another strategy to investigate the cancer initiation dynamics. Because the bulk states correspond to an effective maximum, it is reasonable to assume that the number of pathways going via these states will not be large. This suggests that the dominating pathways on the route to the two-mutations fixation must be those that follow only the boundary states. There are two such pathways, as illustrated in Fig. 3b. Then, instead of trying to account for all possible pathways in the system, the realistic description of the cancer initiation process can be obtained by considering only these two pathways. This significantly simplifies theoretical analysis, and it also allows to understand better the underlying microscopic picture.

The dynamics of cancer initiation in the stochastic model that accounts only for the boundary states (see Fig. 3b) can be explicitly analyzed as shown in detail in the SI. More specifically, the expression for the two-mutations fixation probability starting from the state (1, 0) is given by

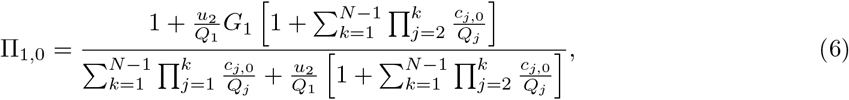

where

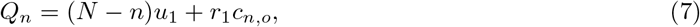

is the forward transition rate (Fig. 3b) for the states on the horizontal axis (*n*_2_ = 0); and

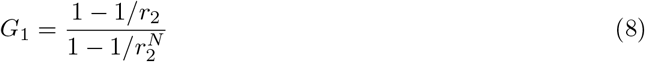

is the fixation probability starting from the state (0, 1) (one double-mutated cell).

The results of theoretical calculations are presented in Fig. 4 for various sets of parameters. It is found that in the weak mutation limit (*u*_1_, *u*_2_ ≪ 1*/N*), the fixation probability depends strongly on the fitness parameter *r*_1_ and weakly on the fitness parameter *r*_2_. This is because in this limit Eq. (6) simplifies into

**Figure 4.**
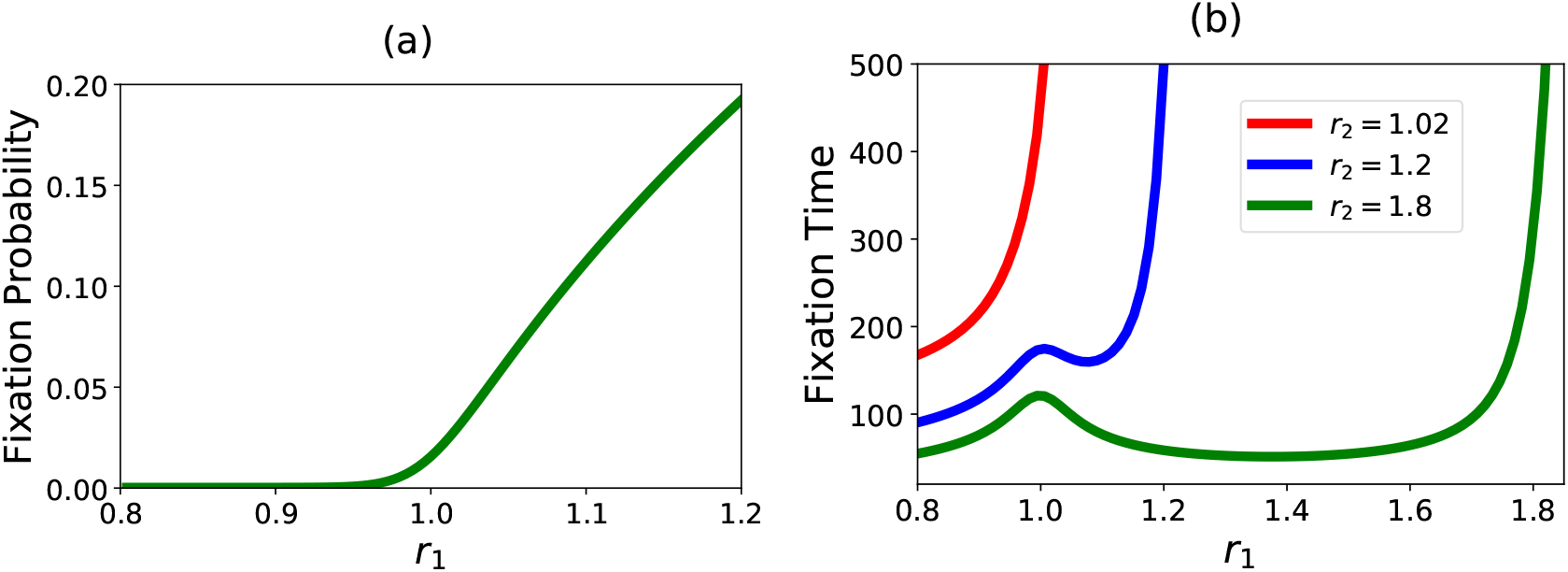
a) Fixation probability for the model with only boundary states as a function of the fitness parameter *r*_1_ for different values of the fitness parameter *r*_2_. Note that several curves are on the top of each other, i.e., no dependence on *r*_2_ is observed. b) Mean fixation times for the model with only boundary states as a function of the fitness parameter *r*_1_ for different values of the fitness parameter *r*_2_. In calculations, the parameters 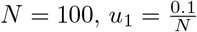 and 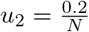 were utilized.

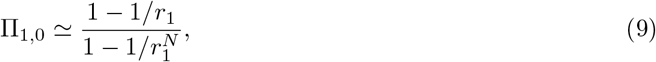

which is essentially the expression for the fixation of the first mutation. One can easily understand this from the scheme in Fig. 3b. In the weak mutation limit, the dominating pathway is the horizontal-diagonal path, while the vertical path has a very low contribution. In addition, it is enough to reach the state (*N*, 0) for evaluating the overall fixation probability because the transition to this state is irreversible. These arguments suggest that the overall fixation of two mutations in the model with only boundary states can be well approximated by the fixation of the first mutation.

The mean fixation times can also be derived using the same method of calculations as for the fixation probabilities as explained in the SI. But it is more instructive to use the following arguments to derive the approximate expression, which also explains better the microscopic processes in the system. Utilizing the previous theoretical analysis [31, 32], the mean fixation time for two mutations can be approximated as

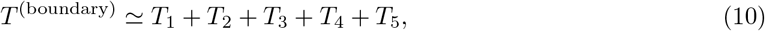

where

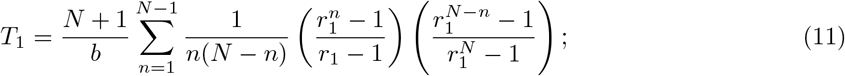

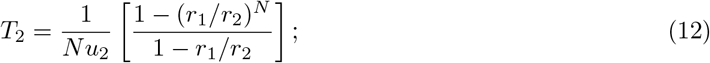

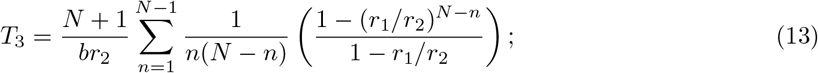

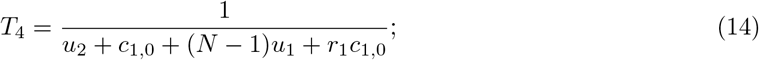

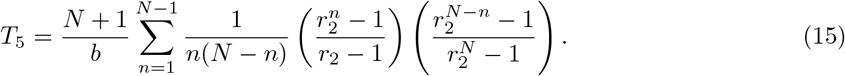

We also have from Eq. (2) that 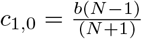. This expression implies that there are five contributions to the mean fixation time. The first term, *T*_1_, describes the time it takes to move along the horizontal pathway to reach the state (*N*, 0), and it corresponds to the mean fixation time of only the first mutation. The second term, *T*_2_, evaluates the average time to move from the horizontal path to the diagonal path. It reflects possible multiple reversible transitions between the state (*N*, 0) and (*N* − 1, 1). The third term, *T*_3_, describes the average time to be on the diagonal path before reaching the full fixation. The fourth term, *T*_4_, is the time to jump from the horizontal path to the vertical path, and it is given by the residence time in the state (1, 0). Finally, the fifth term, *T*_5_, describes the average time on the vertical path before reaching the fixation. Our approximate expression for the mean fixation times has been tested with Monte Carlo computer simulations (not shown here), and good agreement is found, especially in the most biologically relevant regions of *r*_1_ *>* 1 and *r*_2_ *>* 1.

The results of calculations for the mean fixation times in the model with only boundary states is presented in Fig. 4b. Two different behaviors are observed depending on the value of the fitness parameter *r*_2_. If *r*_2_ is close to 1, then the monotonic increase in the overall fixation time as a function of the fitness parameter *r*_1_ is predicted (Fig. 4b). One can understand this by looking into the scheme in Fig. 3b. The system spends most of the time on the diagonal path. Increasing *r*_1_ significantly slows down the motion on this path in the direction of the fixation since the single-mutated cells can replicate faster than the double-mutated cells for *r*_1_ *> r*_2_. In addition, the motion on the vertical path is also not fast due to *r*_2_ ≃ 1. More interesting behavior with a non-monotonic dependence on *r*_1_ is observed for larger values of *r*_2_: see Fig. 4b. This can be again explained by exploring the stochastic scheme in Fig. 3b. While *r*_2_ *> r*_1_, the slowest step in the overall fixation is a translocation along the horizontal path. Clearly, increasing *r*_1_ will decrease this term and accelerate the overall dynamics. However, for *r*_2_ *< r*_1_ the motion along the diagonal path will become slow. These two factors lead to the overall non-monotonic behavior in the system.

We hypothesize that the cancer initiation process can be well described by analyzing only those pathways that follow the boundary states. Our idea is supported by constructing an effective free-energy landscape (Fig. 3a). To fully test this proposal, we analyze the dynamics of fixation of two mutations in the full model (with all possible discrete states) and in the model with only boundary states using Monte Carlo computer simulations for realistic sets of parameters. The results are presented in Fig. 5. In order to fairly compare the dynamics of cancer initiation, both models (all states and only boundary states) have been studied only with the computer simulations, although our analytical calculations, as explained above, agree with the computer simulations predictions for the model with only boundary states.

**Figure 5.**
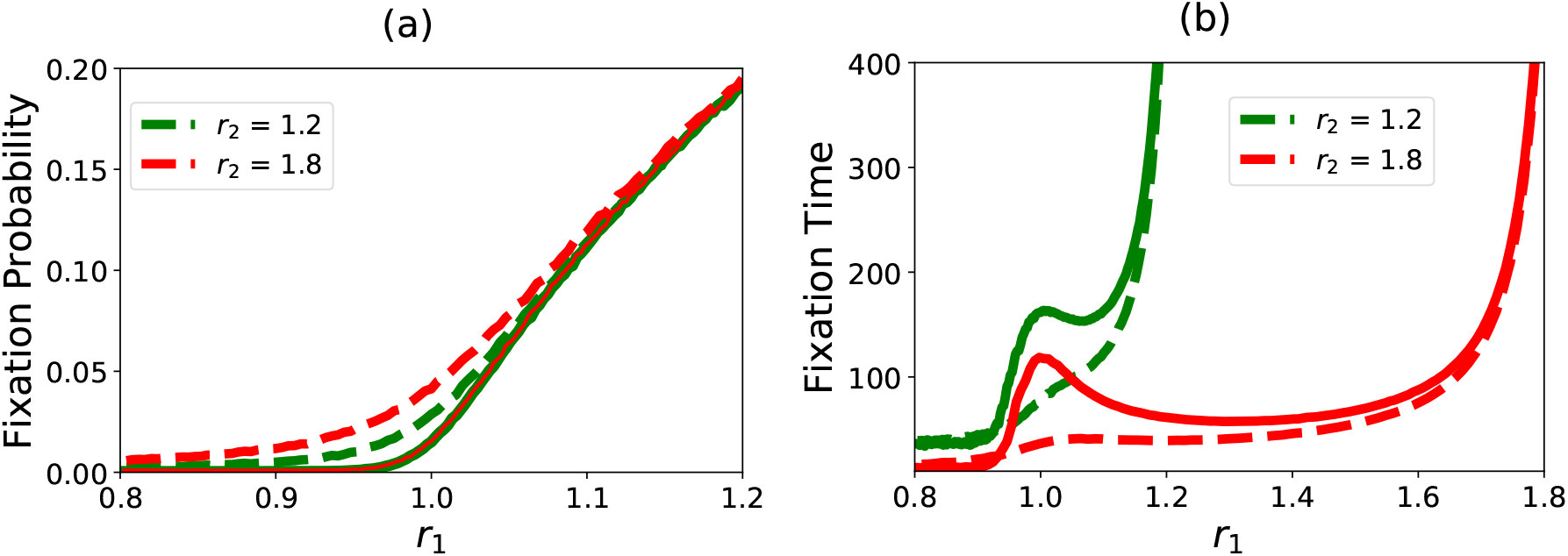
a) Fixation probability for the full model versus the model with only boundary states. b) Mean fixation times for the full model versus the model with only boundary states. Solid lines corresponds to the model with only boundary states, while dashed curves describe the full model. In calculations, the parameters 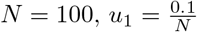 and 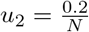 were utilized.

The comparison of fixation probabilities is presented in Fig. 5a, showing a very good agreement for all ranges of parameters. Slight deviations between the model with optimal pathways and the full model are observed only near *r*_1_ ≃ 1. The advantage of our theoretical picture is that these observations can be explained using the schemes in Fig. 2a and Fig. 3b. Due to weak mutations rates (*u*_1_, *u*_2_ ≪ 1*/N*), for *r*_1_ ≃1 the system spends a significant amount of time in the horizontal axis of states with *n*_2_ = 0. This opens the possibility of stochastic tunneling, i.e., moving to the bulk states and crossing them on the road to the overall fixation. Increasing the fitness parameter *r*_1_ moves the system faster to the diagonal axis of states, from which the stochastic tunneling is not possible: see Fig. 3b.

The mean fixation times for the full model and for the model with only boundary states are presented in Fig. 5b. While the deviations in the fixation times are larger than the differences in the fixation probabilities in some regions, the overall agreement is quite satisfactory. The model with the optimal pathways correctly describes the overall trends and quantitatively agrees with the full model for *r*_1_ *<* 1 and for large values of this fitness parameter. The difference found at the region around *r*_1_ ≃ 1 can be explained using the same arguments as above. In this region, the system is originally found mostly on the horizontal axis (see Fig. 3b), opening the possibilities for exploring the side-pathways via the bulk states (stochastic tunneling). But from these pathways the system can escape to the vertical and diagonal axis faster than in the optimal-pathway model. This explains why the mean fixation times in the model with the boundary states in the region around *r*_1_ ≃ 1 are larger than the mean fixation times in the full model.

## Discussions

Although there are some deviations between the predictions of the optimal-pathway model and the full model of two-mutations fixation, it can be argued that a very simple theoretical picture based on utilizing the effective free-energy landscape is capturing many features in the cancer initiation dynamics. Importantly, it allows to significantly simplify the analysis of the complex processes associated with the tumor formation. Instead of looking at very large number of possible pathways on the road to fixation (visiting ∼ *N* ^2^ states), one might concentrate on the processes that are going via very few pathways (visiting ∼ *N* states). This allows us to obtain analytical estimates that significantly improve our understanding of the microscopic mechanisms of underlying processes. For example, the order of mutations [32] and the role of the fitness parameters now can be explained much better.

This theoretical approach can be easily generalized to the system with more than two mutations. If the tumor appears after the fixation of *M* mutations, then the overall number of states in the system will be ∼*N*^*M*^, and it will be prohibitively difficult to analyze such systems even with the most advanced computational tools. Our method suggests that, instead, one could mostly account only for ∼ *N*^*M−*1^ boundary states. One could visualize these processes by noting that the overall space of discrete states is occupying the diagonally cut half of the *M* -dimensional cube, and the boundary states correspond to the *M* + 1 surfaces surrounding this figure. Furthermore, our theoretical results suggest that in the cancer initiation processes the dominating pathways are those that are less heterogeneous from the point of view of different types of cells. In other words, in the system with *M* mutations there are *M* + 1 types of cells that might exists simultaneously (where the number of mutations range from zero to *M*), and we predict that the road to fixation follows mostly via the states with the smaller number of types of cells. From microscopic point of view, from the more heterogeneous states there are more ways of escaping to less heterogeneous states. This is another way of looking at the effective free-energy landscape of the cancer initiation process.

Another advantage of the proposed theoretical method is that it can be sequentially improved by again using the effective free-energy landscape in Fig. 3a. To make the description of the dynamic properties of the fixation process more precise, one might add few more states to the boundary states. These additional states lie near the corners of the stochastic scheme in Fig. 3a. It is expected that this will make the predictions for the fixation probabilities and mean fixation times much closer to the exact values. But importantly, this should not significantly complicate the calculations of dynamic properties.

Although our theoretical approach successfully captures the main physical features of the cancer initiation process with multiple mutations, it is important to discuss its limitations and future directions. Because before the formation of the tumor the total number of tissue cells is constant, the cell replications must be accompanied by cell deaths. Following the typical procedure adopted in the field [22], we utilized the Moran process, which is a mathematical procedure that allows to keep the total number of cells in the system fixed by following specific rules. This procedures assumes that after one cell divides any other cell in the system can be instantly removed with the same probability, which effectively neglects the spatial structure of the tissue. But taking into account the tissue structure, for example, by utilizing the Moran process on graphs [18], might lead to very complex dynamic phenomena. This direction will be interesting to explore. However, there might be a more general problem since the exact biochemical mechanisms of how the normal tissue is regulating cell replications and deaths is still not well understood [19]. In other words, the use of the Moran process might not be a reasonable approach, potentially leading to some artifacts in the theoretical analysis of the cancer initiation dynamics. One of the possible future directions is to utilize more mechanical approaches that utilize various ideas from chemistry and physics in clarifying the cell replications and cell deaths processes [29]. It will be important to explore various theoretical and experimental tools for for more quantitative investigations of cancer initiation processes.

## Author Contributions

H.T. and A.B.K. designed the research. H.T. and C.S carried out the research. H.T. and A.B.K. wrote the article.

## Acknowledgments

We acknowledge the support from the Welch Foundation (C-1559), from the NSF (CHE-1953453 and MCB-1941106), and from the Center for Theoretical Biological Physics sponsored by the NSF (PHY-2019745).

