## Supporting information for "Optimal pathways control fixation of multiple mutations during cancer initiation"

February 7, 2022

In this supporting information we provide details of calculations for the equations in the main text.

#### 1 Calculation of Auxiliary Transition Rates

In this section, we compute the transition rates $a_{n_1,n_2}$, $c_{n_1,n_2}$ and $d_{n_1,n_2}$ that are important for investigating the dynamics of the system. First, we calculate the transition rate $c_{n_1,n_2}$, which describes hopping from state $n_1, n_2$ to state $n_1+1, n_2$. Following calculations presented in Ref. [1], we consider the cell division as a two-state process, as shown in Fig. S1a. Thus, the mean first-passage time from state $(n_1, n_2)$ to the state $(n_1+1, n_2)$ is given by:

$$T_{n_1,n_2,n_1+1,n_2} = \frac{r_1 b n_1 + A(n_1+1) + A(N-n_1-n_2)}{r_1 b n_1 A(N-n_1-n_2)} = \frac{r_1 b n_1 + A(N-n_2+1)}{r_1 b n_1 A(N-n_1-n_2)}, \quad \text{(S1)}$$

which is equal to the inverse transition rate between these states. For $A >> b$ that corresponds to the fact the the cell removal must be instantaneous, we obtain

$$c_{n_1,n_2} = \frac{1}{T_{n_1,n_2,n_1+1,n_2}} \simeq \frac{b n_1 (N-n_1-n_2)}{N-n_2+1}. \quad \text{(S2)}$$

Similarly for the scheme in Fig. S1b we obtain the mean first-passage time from state $(n_1, n_2)$ to the state $(n_1-1, n_2+1)$ is given by

$$T_{n_1,n_2,n_1-1,n_2+1} = \frac{A n_1 + r_2 b n_2 + A(n_2+1)}{r_2 b n_2 A n_1} = \frac{r_2 b n_2 + A(n_1+n_2+1)}{r_2 b n_2 A n_1}. \quad \text{(S3)}$$

Again, for $A >> b$ it reduces to

$$d_{n_1,n_2} = \frac{1}{T_{n_1,n_2,n_1-1,n_2+1}} \simeq \frac{b n_1 n_2}{n_1+n_2+1} \quad \text{(S4)}$$

Finally, for the scheme in Fig. S1c we obtain the mean first-passage time from the state $(n_1, n_2)$ to the state $(n_1, n_2+1)$ is equal to

$$T_{n_1,n_2,n_1,n_2+1} = \frac{A(n_2+1) + r_2 b n_2 + A(N-n_2-n_1)}{r_2 b n_2 A(N-n_1 n_2)} = \frac{r_2 b n_2 + A(N-n_1+1)}{r_2 b n_2 A(N-n_1 n_2)}. \quad \text{(S5)}$$

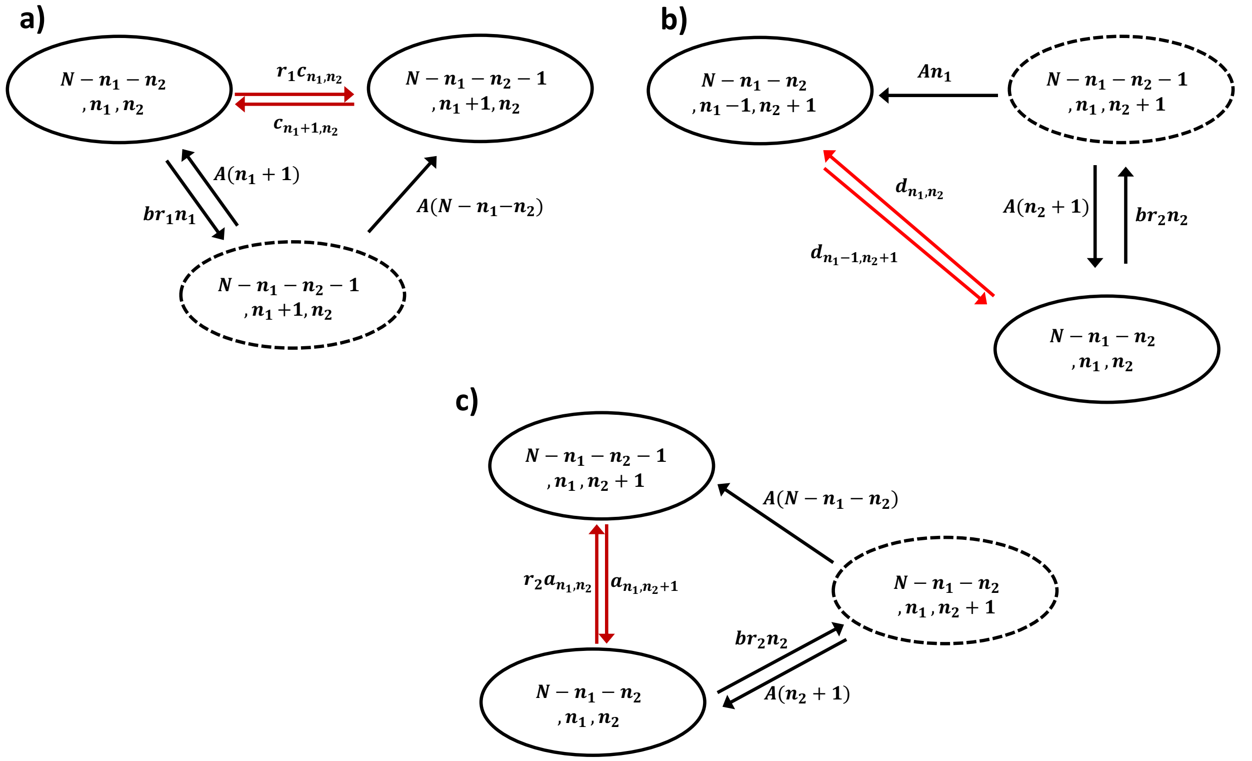


Figure 1: Schematic view for the derivation of transition rates. a) $c_{n_1,n_2}$. b) $d_{n_1,n_2}$. c) $a_{n_1,n_2}$.

For $A >> b$, we derive,

$$a_{n_1,n_2} = \frac{1}{T_{n_1,n_2,n_1,n_2+1}} \simeq \frac{bn_2(N - n_1 - n_2)}{N - n_1 + 1}; \tag{S6}$$

#### 2 Full Analytic Solution for $N = 2$

In this section, we obtain a full exact solution for the first passage probability density functions $F(n_1, n_2; t)$ for $N = 2$ (see Fig. S2). In this case, there are four probability functions, which, for simplicity, we represent as $F_{11}$, $F_{20}$, $F_{10}$, $F_{01}$. The temporal evolution of these functions are governed by the corresponding backward master equations,

$$\frac{dF_{20}}{dt} = -2u_2 F_{20}(t) + 2u_2 F_{11}(t) \tag{S7}$$

$$\frac{dF_{10}}{dt} = -(u_2 + u_1 + r_1 c_1 + c_1)F_{10}(t) + u_2 F_{01}(t) + (u_1 + r_1 c_1)F_{20}(t) \tag{S8}$$

$$\frac{dF_{11}}{dt} = -(u_2 + r_2 d_1 + r_1 d_1)F_{20}(t) + (u_2 + r_2 d_1)F_{02}(t) + r_1 d_1 F_{20}(t) \tag{S9}$$

$$\frac{dF_{01}}{dt} = -(u_1 + r_2 a_1 + a_1)F_{01}(t) + u_1 F_{11}(t) + (r_2 a_1)F_{02}(t) \tag{S10}$$

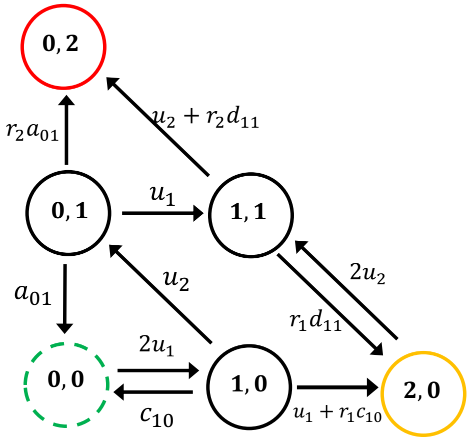


Figure 2: A schematic view of the all possible pathways associated to the fixation of two mutations for $N = 2$.

Performing Laplace transform, we get

$$-\widetilde{F_{11}}(s)(d_1r_1 + d_1r_2 + s + u_2) + d_1\widetilde{F_{20}}(s)r_1 + d_1r_2 + u_2 = 0 \quad \text{(S11)}$$

$$-\widetilde{F_{20}}(s)(s + 2u_2) + 2\widetilde{F_{11}}(s)u_2 = 0 \quad \text{(S12)}$$

$$-\widetilde{F_{01}}(s)(a_1r_2 + a_1 + s + u_1) + a_1r_2 + \widetilde{F_{11}}(s)u_1 = 0 \quad \text{(S13)}$$

$$-\widetilde{F_{10}}(s)(c_1r_1 + c_1 + s + u_1 + u_2) + \widetilde{F_{20}}(s)(c_1r_1 + u_1) + \widetilde{F_{10}}(s)u_2 = 0 \quad \text{(S14)}$$

After some algebra, it can be shown that

$$\widetilde{F_{10}}(s) = \frac{\Phi}{(a_1r_2 + a_1 + s + u_1)(c_1r_1 + c_1 + s + u_1 + u_2)(d_1r_1s + d_1r_2s + 2d_1r_2u_2 + s^2 + 3su_2 + 2u_2^2)} \quad \text{(S15)}$$

where an auxiliary function $\Phi$ is defined as

$$\begin{aligned}\Phi = {} & -2a_1c_1d_1r_1r_2^2u_2 - 2a_1c_1d_1r_1r_2u_2 - 2a_1c_1r_1u_2^2 - 2a_1c_1r_1r_2u_2^2 \\ & -a_1d_1r_2^2su_2 - a_1d_1r_1r_2su_2 - 2a_1d_1r_2^2u_2^2 - 2a_1d_1r_2^2u_1u_2 \\ & -2a_1d_1r_2u_1u_2 - a_1r_2s^2u_2 - 3a_1r_2su_2^2 - 2a_1r_2u_2^3 \\ & -2a_1r_2u_1u_2^2 - 2a_1u_1u_2^2 - 2c_1d_1r_1r_2su_2 - 2c_1d_1r_1r_2u_1u_2 - 2c_1r_1su_2^2 \\ & -2c_1r_1u_1u_2^2 - 3d_1r_2su_1u_2 - 2d_1r_2u_1u_2^2 - 2d_1r_2u_1^2u_2 - 3su_1u_2^2 - 2u_1u_2^3 - 2u_1^2u_2^2\end{aligned} \quad \text{(S16)}$$

Expanding this function in terms of variable $s$ yields the fixation probability,

$$\Pi_{10} = \widetilde{F_{10}}(s = 0) = \frac{a_1c_1r_1 + a_1c_1r_1r_2 + a_1r_2u_1 + a_1r_2u_2 + a_1u_1 + c_1r_1u_1 + u_1^2 + u_2u_1}{(a_1r_2 + a_1 + u_1)(c_1r_1 + c_1 + u_1 + u_2)} \quad \text{(S17)}$$

and, the fixation time,

$$T_{10} = \frac{-\frac{\partial \widetilde{F_{10}}}{\partial s}|_{s=0}}{\widetilde{F_{10}}(s = 0)} = -\frac{\Lambda}{\Pi_{10}} \quad \text{(S18)}$$

where a parameter $\Lambda$ is given by

$$\Lambda = -\frac{(d_1r_1 + d_1r_2 + 3u_2)(a_1c_1r_1 + a_1c_1r_1r_2 + a_1r_2u_1 + a_1r_2u_2 + a_1u_1 + c_1r_1u_1 + u_1^2 + u_2u_1)}{2u_2(a_1r_2 + a_1 + u_1)(c_1r_1 + c_1 + u_1 + u_2)(d_1r_2 + u_2)} + \frac{a_1d_1r_2^2 + a_1d_1r_1r_2 + 3a_1r_2u_2 + 2c_1d_1r_1r_2 + 2c_1r_1u_2 + 3d_1r_2u_1 + 3u_1u_2}{2(a_1r_2 + a_1 + u_1)(c_1r_1 + c_1 + u_1 + u_2)(d_1r_2 + u_2)} + \frac{-a_1c_1r_1 - a_1c_1r_1r_2 - a_1r_2u_1 - a_1r_2u_2 - a_1u_1 - c_1r_1u_1 - u_1^2 - u_2u_1}{(a_1r_2 + a_1 + u_1)^2(c_1r_1 + c_1 + u_1 + u_2)} + \frac{-a_1c_1r_1 - a_1c_1r_1r_2 - a_1r_2u_1 - a_1r_2u_2 - a_1u_1 - c_1r_1u_1 - u_1^2 - u_2u_1}{(a_1r_2 + a_1 + u_1)(c_1r_1 + c_1 + u_1 + u_2)^2} \quad \text{(S19)}$$

In Fig. S3, we compared the analytical results for the fixation probability and mean fixation times with Monte-Carlo computer simulations. Excellent agreement is found for all ranges of parameters.

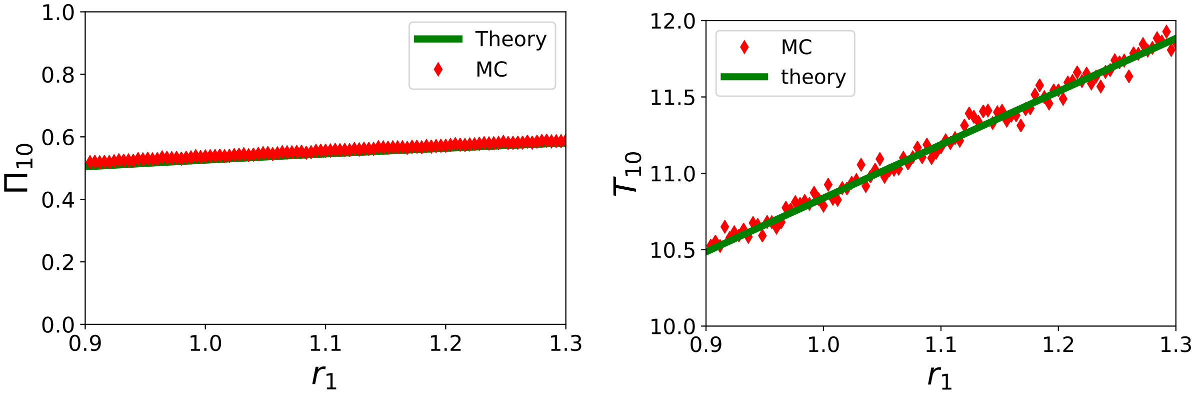


Figure 3: Comparison of computer simulations and exact dynamic properties (fixation probability and mean fixation time) for $N = 2$ case.

#### 3 Full Analytic Solution for $N = 3$

For $N = 3$, there are eight probability functions, which we represents as $F_{11}$, $F_{20}$, $F_{10}$, $F_{01}$, $F_{21}$, $F_{30}$, $F_{02}$, $F_{12}$. The schematic picture of the model is shown in Fig. S4. The corresponding

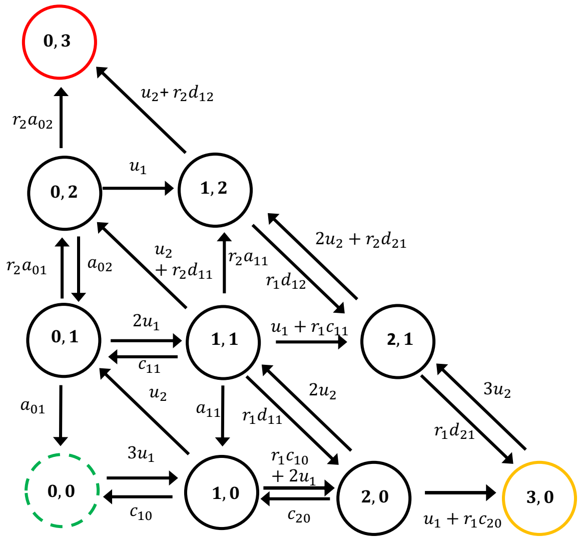


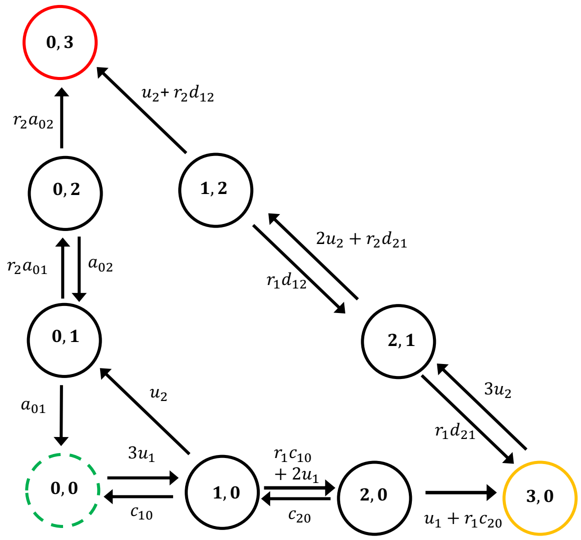


Figure 4: A schematic view of the all possible pathways associated to the fixation of two mutation for $N = 3$: (left) The full model with internal state, and (right) The model with boundary states.

backward master equations for temporal evolution of first-passage probabilities read as

$$\frac{dF_{20}}{dt} = -(2u_2 + c_{20} + u_1 + r_1c_{20})F_{20}(t) + 2u_2F_{11}(t) + c_{20}F_{10}(t) + u_1 + r_1c_{20}F_{30}(t) \quad \text{(S20)}$$

$$\frac{dF_{11}}{dt} = -(a_{11} + r_1d_{11} + u_1 + r_1c_{11} + r_2a_{11} + u_2 + r_2d_{11} + c_{11})F_{11}(t) + a_{11}F_{10}(t)$$
$$+r_1d_{11}F_{20}(t) + (u_1 + r_1c_{11})F_{21}(s) + r_2a_{11}F_{12}(t) + (u_2 + r_2d_{11})F_{02}(t) + c_{11}F_{01}(t) \quad \text{(S21)}$$

$$\frac{dF_{10}}{dt} = -(u_2 + c_{10} + 2u_1 + r_1c_{10})F_{10}(t) + u_2F_{01}(t) + 2u_1 + r_1c_{10}F_{20}(t) \quad \text{(S22)}$$

$$\frac{dF_{01}}{dt} = -(2u_1 + a_{01} + r_2a_{01})F_{01}(t) + 2u_1F_{11}(t) + r_2a_{01}F_{02}(t) \quad \text{(S23)}$$

$$\frac{dF_{21}}{dt} = -(2u_2 + r_2d_{21} + r_1d_{21})F_{21}(t) + r_1d_{21}F_{30}(t) + 2u_2 + r_2d_{21}F_{12}(t) \quad \text{(S24)}$$

$$\frac{dF_{30}}{dt} = -(3u_2)F_{30}(t) + 3u_2F_{21}(t) \quad \text{(S25)}$$

$$\frac{dF_{02}}{dt} = -(u_1 + r_2a_{02} + a_{02})F_{02}(t) + r_2a_{02}F_{03}(t) + a_{02}F_{01}(t) + u_1F_{12}(t) \quad \text{(S26)}$$

$$\frac{dF_{12}}{dt} = -(r_1d_{12} + u_2 + r_2d_{12})F_{12}(t) + r_1d_{12}F_{21}(t) + u_2 + r_2d_{12}F_{03}(t) \quad \text{(S27)}$$

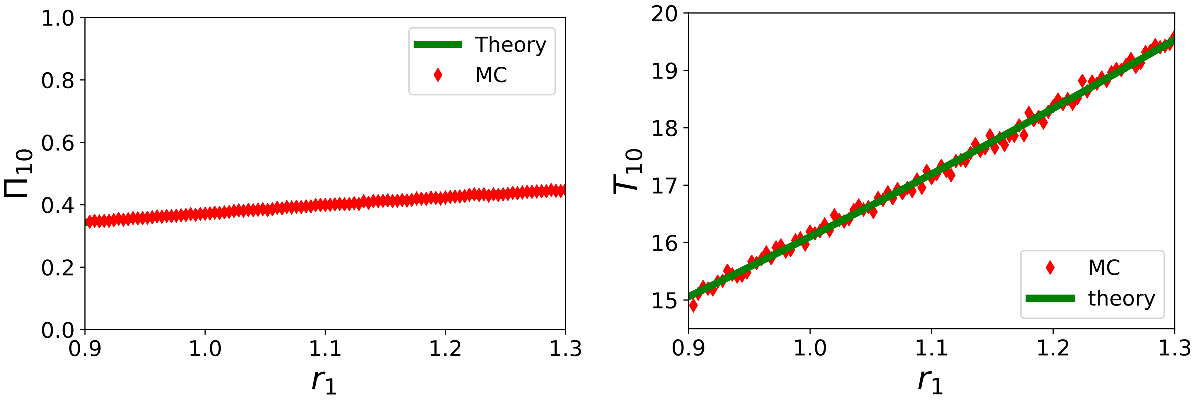


Figure 5: Comparison of computer simulations and exact solution for $N = 3$ (full model).

Performing Laplace transform, one derives

$$(s + 2u_2 + c_{20} + u_1 + r_1c_{20})\widetilde{F_{20}}(s) = 2u_2\widetilde{F_{11}}(s) + c_{20}\widetilde{F_{10}}(s) + u_1 + r_1c_{20}\widetilde{F_{30}}(s) \tag{S28}$$

$$\widetilde{F_{11}}(s)(s + a_{11} + r_1d_{11} + u_1 + r_1c_{11} + r_2a_{11} + u_2 + r_2d_{11} + c_{11}) = a_{11}\widetilde{F_{10}}(s)$$
$$+r_1d_{11}\widetilde{F_{20}}(s) + (u_1 + r_1c_{11})\widetilde{F_{21}}(s) + r_2a_{11}\widetilde{F_{12}}(s) + (u_2 + r_2d_{11})\widetilde{F_{02}}(s) + c_{11}\widetilde{F_{01}}(s) \tag{S29}$$

$$(s + u_2 + c_{10} + 2u_1 + r_1c_{10})\widetilde{F_{10}}(s) = u_2\widetilde{F_{01}}(s) + 2u_1 + r_1c_{10}\widetilde{F_{20}}(s) \tag{S30}$$

$$(s + 2u_1 + a_{01} + r_2a_{01})\widetilde{F_{01}}(s) = 2u_1\widetilde{F_{11}}(s) + r_2a_{01}\widetilde{F_{02}}(s) \tag{S31}$$

$$(s + 2u_2 + r_2d_{21} + r_1d_{21})\widetilde{F_{21}}(s) = r_1d_{21}\widetilde{F_{30}}(s) + 2u_2 + r_2d_{21}\widetilde{F_{12}}(s) \tag{S32}$$

$$(s + 3u_2)\widetilde{F_{30}}(s) = 3u_2\widetilde{F_{21}}(s) \tag{S33}$$

$$(s + u_1 + r_2a_{02} + a_{02})\widetilde{F_{02}}(s) = r_2a_{02} + a_{02}\widetilde{F_{01}}(s) + u_1\widetilde{F_{12}}(s) \tag{S34}$$

$$(s + r_1d_{12} + u_2 + r_2d_{12})\widetilde{F_{12}}(s) = r_1d_{12}\widetilde{F_{21}}(s) + u_2 + r_2d_{12} \tag{S35}$$

Again, expanding $\widetilde{F_{10}}(s)$ in terms of the variable $s$ yields the fixation probability,

$$\Pi_{10} = \widetilde{F_{10}}(s = 0) \tag{S36}$$

and, the mean fixation time,

$$T_{10} = \frac{-\frac{\partial \widetilde{F_{10}}}{\partial s}|_{s=0}}{\widetilde{F_{10}}(s = 0)}. \tag{S37}$$

Since these expressions are extremely long and bulky, it is not feasible to present them in this supporting information. The corresponding Mathematica file is available. In Fig. S5 we compared these results with Monte-Carlo computer simulations, and excellent agreement is found.

Now we analyze the model for $N = 3$ but with only the boundary states (Fig. S4). In this case, there are only seven states. The corresponding first-passage probability density

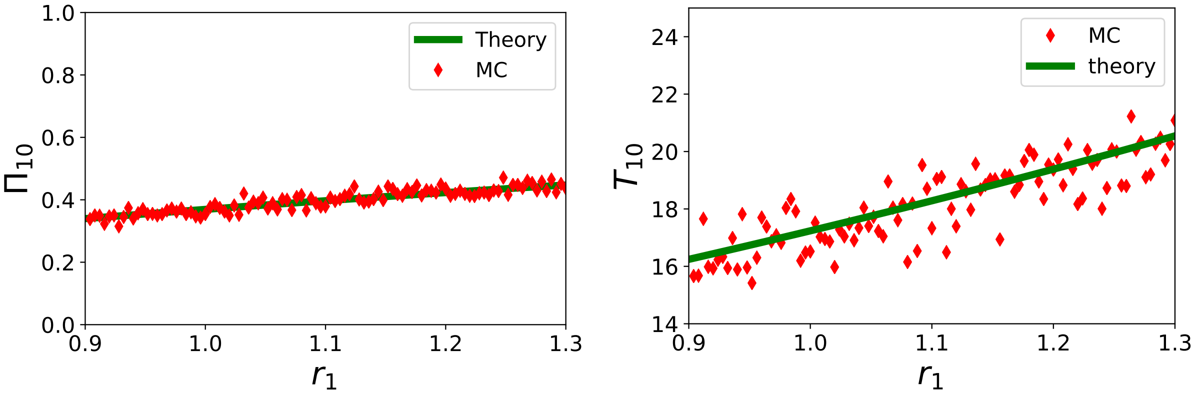


Figure 6: Comparison of computer simulations and exact solution for the model with boundary states ($N = 3$).

functions are governed by following backward master equations,

$$\frac{dF_{20}}{dt} = -(c_{20} + u_1 + r_1 c_{20})F_{20}(t) + c_{20}F_{10}(t) + u_1 + r_1 c_{20}F_{30}(t) \tag{S38}$$

$$\frac{dF_{10}}{dt} = -(u_2 + c_{10} + 2u_1 + r_1 c_{10})F_{10}(t) + u_2 F_{01}(t) + 2u_1 + r_1 c_{10}F_{20}(t) \tag{S39}$$

$$\frac{dF_{01}}{dt} = -(a_{01} + r_2 a_{01})F_{01}(t)r_2 a_{01}F_{02}(t) \tag{S40}$$

$$\frac{dF_{21}}{dt} = -(2u_2 + r_2 d_{21} + r_1 d_{21})F_{21}(t) + r_1 d_{21}F_{30}(t) + 2u_2 + r_2 d_{21}F_{12}(t) \tag{S41}$$

$$\frac{dF_{30}}{dt} = -(3u_2)F_{30}(t) + 3u_2 F_{21}(t) \tag{S42}$$

$$\frac{dF_{02}}{dt} = -(u_1 + r_2 a_{02} + a_{02})F_{02}(t) + r_2 a_{02}F_{03}(t) + a_{02}F_{01}(t) + u_1 F_{12}(t) \tag{S43}$$

$$\frac{dF_{12}}{dt} = -(r_1 d_{12} + u_2 + r_2 d_{12})F_{12}(t) + r_1 d_{12}F_{21}(t) + u_2 + r_2 d_{12}F_{03}(t) \tag{S44}$$

Performing Laplace transformations, we obtain

$$(s + c_{20} + u_1 + r_1 c_{20})\widetilde{F_{20}}(s) = +c_{20}\widetilde{F_{10}}(s) + u_1 + r_1 c_{20}\widetilde{F_{30}}(s) \tag{S45}$$

$$(s + u_2 + c_{10} + 2u_1 + r_1 c_{10})\widetilde{F_{10}}(s) = u_2\widetilde{F_{01}}(s) + 2u_1 + r_1 c_{10}\widetilde{F_{20}}(s) \tag{S46}$$

$$(s + a_{01} + r_2 a_{01})\widetilde{F_{01}}(s) = r_2 a_{01}\widetilde{F_{02}}(s) \tag{S47}$$

$$(s + 2u_2 + r_2 d_{21} + r_1 d_{21})\widetilde{F_{21}}(s) = r_1 d_{21}\widetilde{F_{30}}(s) + 2u_2 + r_2 d_{21}\widetilde{F_{12}}(s) \tag{S48}$$

$$(s + 3u_2)\widetilde{F_{30}}(s) = 3u_2\widetilde{F_{21}}(s) \tag{S49}$$

$$(s + u_1 + r_2 a_{02} + a_{02})\widetilde{F_{02}}(s) = r_2 a_{02} + a_{02}\widetilde{F_{01}}(s) + u_1\widetilde{F_{12}}(s) \tag{S50}$$

$$(s + r_1 d_{12} + u_2 + r_2 d_{12})\widetilde{F_{12}}(s) = r_1 d_{12}\widetilde{F_{21}}(s) + u_2 + r_2 d_{12} \tag{S51}$$

Again, using Eqs. (S36) and (S37) we calculate the fixation probability and mean fixation times. In Fig. S6 we verified these results with Monte Carlo computer simulations.

### 4 Fixation Probabilities for General Model with Boundary States

Let us consider a scheme as presented in Fig. 3b in the main text of the manuscript. Our aim is to calculate the fixation probability for this model. For simplicity in notations, we describe the states on the horizontal branch with $(n, 0)$, which means there are $n$ cells with one mutation and $N-n$ normal cells. Also, all states on the vertical branch are indexed as $(0, n)$, which describes $n$ cells mutated twice and $N-n$ normal cells. We define $F(n, 0; t=0|0, N; t) = F_n(t)$ as first passage probability density to reach the fixation at time $t$ if at $t=0$ the system started in the state $(n, 0)$. The temporal evolution of these probabilities is governed by the following set of backward master equations,

$$\begin{aligned} \frac{dF_1}{dt} &= Q_1 F_2(t) + u_2 G_1(t) - (u_2 + Q_1 + c_1) F_1(t) \\ \frac{dF_2}{dt} &= Q_2 F_3(t) + c_2 F_1(t) - (Q_2 + c_2) F_2(t) \\ \frac{dF_n}{dt} &= Q_n F_{n+1}(t) + c_n F_{n-1}(t) - (Q_n + c_n) F_n(t) \end{aligned} \tag{S52}$$

for $1 \leq n \leq N-2$, and

$$\frac{dF_{N-1}}{dt} = Q_{N-1} F_N(t) + c_{N-1} F_{N-2}(t) - (Q_{N-1} + c_{N-1}) F_{N-1}(t) \tag{S53}$$

for $n = N-1$. In these expressions, we defined $Q_n = (N-n)u_1 + r_1 c_n$ and $c_n = \frac{n(N-n)}{N+1}$. For this system of equations we also have an initial condition, $F_N(t) = \delta(t)$. After performing Laplace transformations, we obtain

$$\begin{aligned} (s + u_2 + Q_1 + c_1)\widetilde{F_1}(s) &= Q_1 \widetilde{F_2}(s) + u_2 \widetilde{G_1}(s) \\ (s + Q_2 + c_2)\widetilde{F_2}(s) &= Q_2 \widetilde{F_3}(s) + c_2 \widetilde{F_1}(s) \\ (s + Q_n + c_n)\widetilde{F_n}(s) &= Q_n \widetilde{F_{n+1}}(s) + c_n \widetilde{F_{n-1}}(s) \\ (s + Q_{N-1} + c_{N-1})\widetilde{F_{N-1}}(s) &= Q_{N-1} + c_{N-1} \widetilde{F_{N-1}}(s) \end{aligned} \tag{S54}$$

Similarly, for the vertical branch, we define $G(0, n; t=0|0, N; t) = G_n(t)$ as a first passage probability density to reach the fixation at time $t$ if at $t=0$ the system started in the state $(0, n)$. Temporal evolution of these functions satisfy the following equations,

$$\begin{aligned} \frac{dG_1}{dt} &= r_2 a_1 G_2(t) - a_2(1 + r_2) G_1(t) \\ \frac{dG_2}{dt} &= r_2 a_2 G_3(t) + a_2 G_1(t) - a_2(1 + r_2) G_2(t) \\ \frac{dG_n}{dt} &= r_2 a_n G_{n+1} + a_n G_{n-1}(t) - a_n(1 + r_2) G_n(t) \end{aligned} \tag{S55}$$

for $1 \leq n \leq N-2$, and

$$\frac{dG_{N-1}}{dt} = r_2 a_{N-1} G_N(t) + a_{N-1} G_{N-2}(t) - a_{N-1}(1 + r_2) G_{N-1}(t) \tag{S56}$$

for $n = N-1$. Here we also have $a_n = \frac{n(N-n)}{N+1}$. For this system of equations, the initial condition is $G_N(t) = \delta(t)$. Applying Laplace transformation yields

$$\begin{aligned}(s + a_1(1+r_2))\widetilde{G_1}(s) &= r_2 a_1 \widetilde{G_2}(s) \\ (s + a_2(1+r_2))\widetilde{G_2}(s) &= r_2 a_2 \widetilde{G_3}(s) - a_2 \widetilde{G_1}(s) \\ (s + a_n(1+r_2))\widetilde{G_n}(s) &= r_2 a_n \widetilde{G_{n+1}}(s) - a_n \widetilde{G_{n-1}}(s) \\ (s + a_{N-1}(1+r_2))\widetilde{G_{N-1}}(s) &= r_2 a_{N-1} \widetilde{G_N}(s) - a_{N-1} \widetilde{G_{N-2}}(s)\end{aligned} \tag{S57}$$

Since we want to calculate only the fixation probabilities, it is convenient to use the following approximation at the steady state conditions ($s \to 0$):

$$\widetilde{G_1}(s) \simeq G_n; \quad \widetilde{F_n}(s) \simeq \Pi_n \tag{S58}$$

Substituting $G_n$ into Eqns S57 yields:

$$\begin{aligned}a_1(1+r_2)G_1 &= r_2 a_1 G_2 \\ a_2(1+r_2)G_2 &= r_2 a_2 G_3 - a_2 G_1 \\ a_n(1+r_2)G_n &= r_2 a_n G_{n+1} - a_n G_{n-1} \\ a_{N-1}(1+r_2)G_{N-1} &= r_2 a_N G_{N+1} - a_N G_{N-1}\end{aligned} \tag{S59}$$

Solving this set of recurrence relation gives $G_1$:

$$G_1 = \frac{1 - \frac{1}{r_2}}{1 - \frac{1}{r_2^N}} \tag{S60}$$

To proceed further, we substitute $\Pi_n$ into Eqs. (S54), leading to

$$\begin{aligned}(u_2 + Q_1 + c_1)\Pi_1 &= Q_1 \Pi_2 + u_2 G_1 \\ (Q_2 + c_2)\Pi_2 &= Q_2 \Pi_3 + c_2 \Pi_1 \\ (Q_n + c_n)\Pi_n &= Q_n \Pi_{n+1} + c_n \Pi_{n-1} \\ (Q_{N-1} + c_{N-1})\Pi_{N-1} &= Q_{N-1} + c_{N-1} \Pi_{N-2}\end{aligned} \tag{S61}$$

Solving these equations produces the final expression for the fixation probability,

$$\Pi_1 = \frac{1 + \frac{u_2}{Q_1} G_1 \left[1 + \sum_{k=1}^{N-1} \prod_{j=2}^{k} \frac{c_{j,0}}{Q_j}\right]}{\sum_{k=1}^{N-1} \prod_{j=1}^{k} \frac{c_{j,0}}{Q_j} + \frac{u_2}{Q_1} \left[1 + \sum_{k=1}^{N-1} \prod_{j=2}^{k} \frac{c_{j,0}}{Q_j}\right]}. \tag{S62}$$

### 5 Dependence on the Number of Stem Cells

In Fig. S7, we plotted the fixation probabilities and the mean fixation time as a function of the number of stem cells in the tissue $N$. One can see a perfect agreement for the fixation probabilities and a semi-quantitative agreement for the mean fixation times.

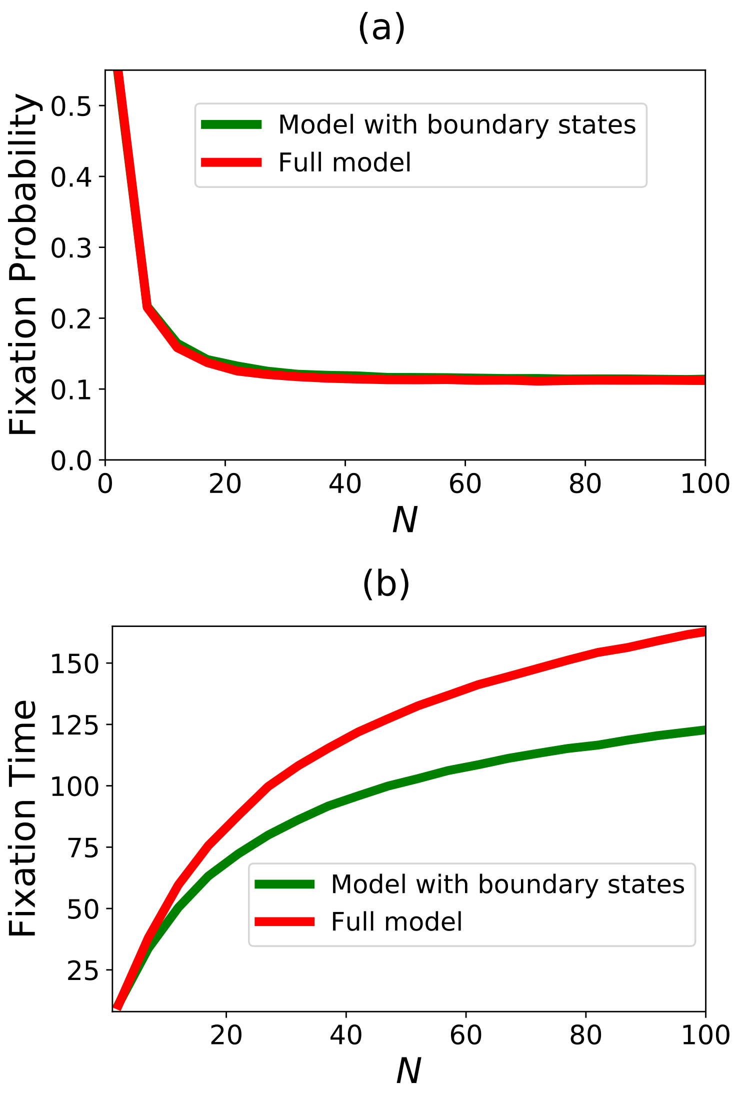


Figure 7: Fixation probabilities and fixation time versus $N$. In calculations, the parameters $N = 100$, $u_1 = \frac{0.1}{N}$, $u_2 = \frac{0.2}{N}$, $r_1 = 1.1$ and $r_2 = 1.2$ were utilized.
